# Origin of plant trait data matters: Shared species of Northwestern Europe and the Pannonian Ecoregion have different trait values in the two regions

**DOI:** 10.1101/2024.10.14.618145

**Authors:** Judit Sonkoly, Péter Török

## Abstract

Trait-based ecology considerably increased our comprehension of various fields related to ecology and evolution. As measuring traits can be time-consuming and costly, analyses regularly gather trait data from databases instead of carrying out new measurements. However, intraspecific trait variability can cause considerable differences between the trait values of different population and regions. Here we evaluated whether intraspecific trait variability causes considerable differences in trait values measured in two different regions of Europe. We tested whether regionally measured trait data from the Pannonian Ecoregion differ from trait data of the same species originating from Northwestern Europe, by comparing data from the Pannonian Database of Plant Traits (PADAPT) and the LEDA Traitbase. We evaluated six traits of the same set of species: thousand-seed mass (TSM), seed bank persistence index (SBPI), leaf area (LA), leaf dry matter content (LDMC), specific leaf area (SLA), and leaf dry mass. We found that trait data from the two databases significantly differed for TSM, SBPI, SLA, and LDMC. Our results indicate that the markedly different climate of the two regions can cause substantial intraspecific trait variation, therefore, the geographical origin of trait data matters in trait-based analyses. The findings corroborate the assumption that regionally measured trait data and building regional databases are essential for reliable regional-scale trait-based studies. We conclude that for studies analysing traits in the Pannonian Ecoregion (and possibly in Easter and Central Europe in general), it is advisable to use PADAPT instead of databases compiling data from regions with markedly different climatic conditions.

## Introduction

Anthropogenic habitat destruction and climate change make it ever more crucial to understand their growing consequences on biodiversity and ecosystem functions (Cardinale *et al*. 2012). Functional traits can provide us with tools to reveal and generalise mechanisms related to the formation and maintenance of biodiversity, and to predict how ecosystem functions and services might be affected by changes in biodiversity (Garnier *et al*. 2007, Lavorel *et al*. 2013). This has led to a growing interest in trait-based ecology (Cernansky 2017), which has considerably increased our knowledge and comprehension of various fields related to ecology and evolution (e.g., Lavorel and Garnier 2002, McGill *et al*. 2006, Cadotte *et al*. 2011). However, measuring traits can be time-consuming and costly (Kattge *et al*. 2011, Cordlandwehr *et al*. 2013), therefore, trait data gathered from trait databases is regularly used instead of carrying out new measurements for every single analysis.

As a result of standardised protocols for trait measurements (Cornelissen *et al* 2003, Pérez-Harguindeguy *et al*. 2016) and the enormous efforts invested in compiling trait data in the last decades, many large plant trait databases have been established to facilitate trait-based analyses. Some global databases compile data for a specific group of traits, for example SID (Seed Information Database, Royal Botanical Gardens Kew 2023) or D3 (Dispersal and Diaspore Database, Hintze *et al*. 2013). However, most databases gather data for a wide range of traits at regional scales, such as BiolFlor for the German flora (Klotz *et al*. 2002), BROT for the flora of the Mediterranean region (Tavşanoğlu and Pausas 2018), PLADIAS for the flora of the Czech Republic (Chytrý *et al*. 2021), or the LEDA Traitbase for the flora of Northwestern Europe (Kleyer *et al*. 2008). The largest plant trait database is available in TRY (Kattge *et al*. 2020), which provides a global coverage for a wide range of traits by integrating several databases and datasets.

Interspecific trait variability is generally considered to be much greater than intraspecific variability, which is a basic tenet of studies using a mean trait value for each species (Violle *et al*. 2012, Cordlandwehr *et al*. 2013), thereby treating intraspecific variability as something that could not conceal broad trends (Shipley *et al*. 2016). The assumption that intraspecific variability is greater than interspecific variability has been corroborated by some studies (Garnier *et al*. 2001, Jung *et al*. 2014), but it has also been shown that intraspecific variability is also not necessarily negligible (e.g., Messier *et al*. 2010, Auger and Shipley 2013, Jung *et al*. 2014). Intraspecific trait variability can be a result of genetic differences, phenotypic plasticity, or a mixture of both (Sandquist and Ehleringer 1997, Albert *et al*. 2011), which can cause considerable differences between the trait values of different populations (Albert *et al*. 2012). Some plant traits show more plasticity than others (Shipley *et al*. 2016), for example, traits related to resource acquisition show great variability (Shipley 2000, Violle *et al*. 2009a), while reproductive traits such as seed mass show less (Violle *et al*. 2009b).

It is quite well-known that interspecific trait differences are correlated with environmental conditions such as climate (Hodkinson *et al*. 1998, Wright and Westoby 2002, Poorter *et al*. 2009) or soil conditions (Maire *et al*. 2015). However, the effects of environmental conditions on intraspecific trait variability (see e.g., Garnier *et al*. 2001, Mokany and Ash 2008, Hudson *et al*. 2011) are much less considered, although these effects are particularly relevant for studies using data gathered from databases covering large geographical scales (Cordlandwehr *et al*. 2013). Based on the above considerations, if relevant environmental conditions significantly differ between regions, trait data measured in other regions may differ from the trait values actually representative of the populations in the study area. Methodological decisions such as what database to use and how to clean the available data in order to have a reliable dataset can substantially change the outcome of the analysis (see Augustine *et al*. 2024). This also means that although data for many traits can be relatively easily gathered from large-scale databases, the fact that they include data from multiple regions and often from markedly different environmental conditions compared to the study region may question the applicability of large-scale databases for regional-scale studies.

Our aim was to evaluate whether intraspecific trait variability causes considerable differences in trait values measured in two regions of Europe characterised by markedly different climatic conditions. To this end, we tested whether regionally measured trait data from the Pannonian Ecoregion differ from trait data of the same species originating from Northwestern Europe, by comparing data on six traits from the Pannonian Database of Plant Traits (PADAPT, Sonkoly *et al*. 2023) with data from the LEDA Traitbase (Kleyer *et al*. 2008).

## Materials & Methods

We used the checklist of PADAPT, based which we created paired samples, i.e., we compared trait data from PADAPT and LEDA for the same set of species. We selected those numeric traits for the comparisons for which PADAPT provides a compilation of recent measurements, i.e., thousand-seed mass (TSM), seed bank persistence index (SBPI), leaf area (LA), leaf dry matter content (LDMC), specific leaf area (SLA), and leaf dry mass. We calculated the mean value of all records from PADAPT for each species (Appendix 1). SBPI was calculated following the Seed Longevity Index of Bekker et al. (1998): we express the ratio of data indicating a persistent soil seed bank with a value from 0 to 1, where zero means that all available data indicate a transient seed bank, and 1 means that all available data indicate a persistent seed bank.

Data files for the following traits were obtained from LEDA on 23th November 2021: (i) Seed mass (corresponding to TSM in PADAPT), (ii) Seed longevity (corresponding to SBPI, we calculated SBPI values for each species based on seed longevity data from LEDA), (iii) Leaf size (corresponding to LA), (iv) SLA, (v) LDMC, and (vi) Leaf mass (corresponding to leaf dry mass after converting mg to g). We extracted data from LEDA data files for the species also present in our database and calculated the mean value of all records from LEDA for each species (Appendix 1). To maximise comparability, we considered only those records that were described as ‘actual measurement’. In the case of leaf traits, we excluded measurements without petiole and rachis, and in the case of seed mass, we excluded measurements of multi-seeded generative dispersules. We then compared the data using paired Wilcoxon signed-rank tests in R version 4.0.3 (R Core Team 2020). We decided not to compare our data with data from TRY (Kattge *et al*. 2020) as some of the leaf trait records included in PADAPT 1.0 are also included in TRY.

## Results

Compared to LEDA, PADAPT 1.0 provides thousand-seed mass (TSM) data for further 595 species, seed bank persistence index (SBPI) for further 63 species, leaf area (LA) for further 1,087 species, leaf dry matter content (LDMC) for further 1,098 species, specific leaf area (SLA) for further 616 species, and leaf dry mass (LDM) data for further 1,027 species. TSM, SBPI, SLA, and LDMC values in PADAPT significantly differed from those in LEDA (Table 1, Fig. 1A, 1B, 1D and 1E, respectively), while LA and LDM values did not differ significantly (Table 1, Fig. 1C and 1F, respectively).

**Table 1.** Comparison of data from the LEDA Traitbase and the Pannonian Database of Plant Traits (PADAPT), pairwise Wilcoxon signed-rank tests.

|  | LEDA<br>(mean±SE) | PADAPT<br>(mean±SE) | *p* value | n |
| --- | --- | --- | --- | --- |
| Thousand-seed mass (TSM, g) | 18.79±5.06 | 16.19±4.60 | <0.001*** | 1246 |
| Seed bank persistence index (SBPI) | 0.35±0.01 | 0.68±0.02 | <0.001*** | 468 |
| Leaf area (LA, mm<sup>2</sup>) | 2603.01±261.24 | 2235.51± 220.74 | 0.101 | 494 |
| Specific leaf area (SLA, mm<sup>2</sup>/mg) | 24.92±0.39 | 24.49± 0.47 | <0.001*** | 960 |
| Leaf dry matter content (LDMC, mg/g) | 201.48±3.75 | 251.580± 4.59 | <0.001*** | 475 |
| Leaf dry mass (LDM, mg) | 146.77± 18.94 | 132.59± 17.49 | 0.215 | 490 |

**Figure 1.**
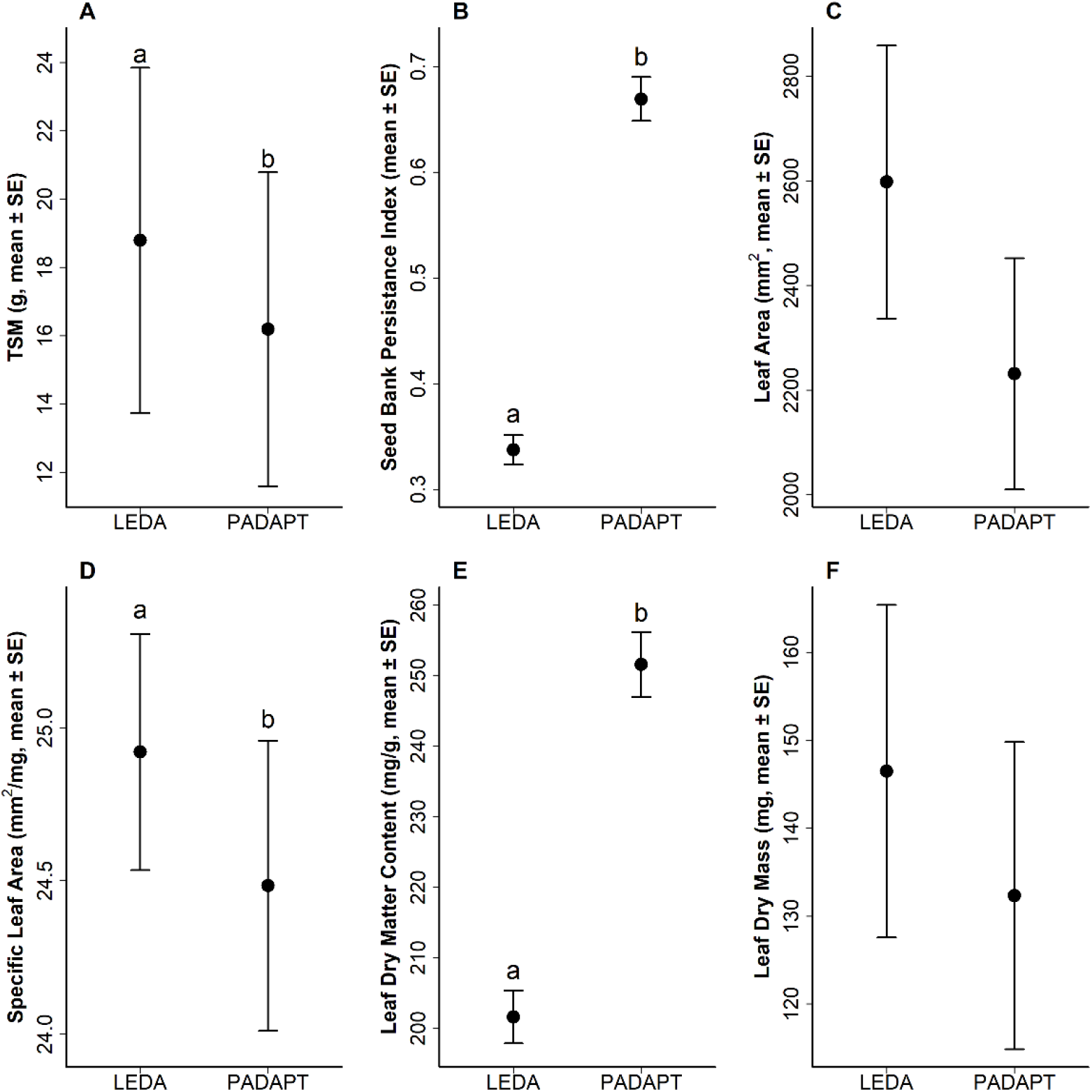
Comparison of data from the LEDA Traitbase and PADAPT for (A) thousand-seed mass (TSWM), (B) seed bank persistence index, (C) leaf area, (D) specific leaf area, (E) leaf dry matter content and (F) leaf dry mass. Dots represent mean values, error bars represent standard error. Wilcoxon signed-rank tests, significant differences are denoted by different letters.

## Discussion

We found that shared plant species of Northwestern Europe and the Pannonian Ecoregion have significantly different values for several traits based on data originating from Northwestern Europe (LEDA Traitbase, Kleyer *et al*. 2008) versus from the Pannonian Ecoregion (PADAPT, Sonkoly *et al*. 2023). These results underline that intraspecific trait variation between regions can be considerable at least for some traits (Albert *et al*. 2011, 2012), and thus the geographical origin of trait data matters in trait-based analyses.

Our finding that thousand-seed mass (TSM) values from PADAPT are significantly lower than TSM values for the same species from the LEDA Traitbase indicates that species of the Pannonian Ecoregion have lower seed mass in this region than the same species in Northwestern Europe (which is the focal region of the LEDA Traitbase). Some results suggest that there is a negative relationship between precipitation and seed mass across species (Baker 1972, Wright and Westoby 1999), more recent studies found that seed mass decreases across species with increasing aridity and precipitation variability (Yu *et al*. 2007, Harel *et al*. 2011). As some previous studies have demonstrated that climatic factors can also cause intraspecific variation in seed mass (e.g., Yeşilyurt *et al*. 2017, Ge *et al*. 2020, Love *et al*. 2022), the difference we observed may be a result of the markedly different climate of Northwestern Europe compared to the Pannonian Ecoregion. The Pannonian Ecoregion is characterised by a more continental climate than Northwestern Europe, with lower precipitation and higher summer temperatures (New *et al*. 2002, Peel *et al*. 2007), and longer sunshine duration (e.g., Kothe *et al*. 2013), which factors can have various effects on the seed size of species.

In contrast with our finding, some previous studies suggest that dry and hot summers should favour the production of larger, not smaller seeds. In stressful environments, such as under dry conditions, regeneration success can be increased by producing larger seeds (Muller-Landau 2010). This is in line with some previous findings, for example *Pinus monophyla* trees growing in drier habitats produced larger seeds than specimens growing in more mesic habitats (Vasey *et al*. 2022), and stressful environments were also associated with larger seeds in the case of *Trichloris crinita* (Marinoni *et al*. 2018). The finding that higher annual minimum temperatures were associated with lower seed mass in *Helianthemum salicifolium* (Yeşilyurt *et al*. 2017) is also in contrast with our results. On the other hand, in line with the detected lower seed mass in the Pannonian Ecoregion, Harel *et al*. (2011) found that the seed mass of most of the studied species increased with increasing precipitation of the habitat. Dry conditions experienced by *Desmodium paniculatum* individuals resulted in the production of lighter seeds (Wulff 1986), which is in line with our results. All things considered, it is highly likely that climate plays a vital role in this difference, but exactly what climatic factor plays the most important role in creating the detected lower seed mass in the Pannonian Ecoregion cannot be determined based on our results, warranting detailed studies of the issue.

The considerable difference detected in seed bank persistence index (SBPI) is probably at least partly caused by the average number of individual records per species based on which SBPI was calculated being an order of magnitude greater in LEDA than in PADAPT. Results indicating persistent seed banks are easier to obtain than results indicating transient seed banks, because if a species is missing from the seed bank it does not necessarily mean that the species does not have persistent seeds, mainly because seeds of species with low seed production can easily be missed by soil seed bank sampling (Saatkamp *et al*. 2009). This can result in higher SBPI values (indicating more persistent seeds) when the number of data points per species is low, such as in the case of PADAPT.

However, the observed difference may also be caused by climatic factors to some extent. For example, the seed viability of species of dry habitats decreases when they experience moist conditions, mostly due to pathogenic fungi (Blaney and Kotanen 2001, Schafer and Kotanen 2003). Chen *et al*. (2021) also found that the seed persistence of all the 11 studied species decreased with increasing precipitation, which effect could be attributed to the action of soil fungi. These results can at least partly explain our finding that the studied species had drastically lower seed bank persistence based on data from the more humid climate of Northwestern Europe than based on data from the Pannonian Ecoregion. On the other hand, some other studies suggest that increased soil temperatures can speed up processes leading to the decline of seed viability (Ooi 2012), moreover, parent plants experiencing high temperatures produce less dormant seeds (Fenner 1991, Kochanek *et al*. 2010). In contrast with our results, these effects could lead to less persistent seed banks in habitats experiencing high temperatures. Nevertheless, it seems highly likely that soil seed survival, and ultimately the seed bank persistence of a species can vary greatly between sites and regions (Saatkamp *et al*. 2009).

Studies analysing leaf traits across species have generally found that plant species inhabiting regions with drier climates tend to have thicker leaves with low SLA and high LDMC values (e.g., Fonseca *et al*. 2000, Wright *et al*. 2004), because low SLA is linked with improved efficiency of water-use in case of drought tress (Wellstein *et al*. 2017). Studies of the effects of climate on the intraspecific variation of SLA and LDMC have found a pattern consistent with the one observed across species: lower SLA values were found in plants exposed to lower water availability (Garnier *et al*. 2001, Cornwell and Ackerly 2009, Wellstein *et al*. 2017), which is consistent with other findings about plastic trait changes allowing plants to withstand drought stress (Niinemets 2001, Chaves *et al*. 2002).

Although we did not find a significant difference between the leaf area data of the same species originating from Northwestern Europe versus the Pannonian Ecoregion, some previous studies demonstrated that differences in water availability can result in differences in leaf area for some species, because leaf size is related to water and energy balance (Cornelissen *et al*. 2003). However, the direction of the effect is not clear; some studies found that plants growing under more arid conditions have higher leaf area values (e.g., Sandquist and Ehleringer 1997), but some other results indicate that leaf area increases with increasing water availability (Cornwell and Ackerly 2009). Nevertheless, the fact that there was no significant difference in leaf area and leaf dry mass between the two databases may reflect that leaf size in itself is a less important factor of the leaf economic spectrum than LDMC and SLA (Wright *et al*. 2004), which are more directly related to the rate of photosynthetic activity (e.g., Reich *et al*. 1998, Reich 2014).

In conclusion, our results indicate that climatic differences between regions can cause substantial intraspecific trait variation, therefor, the geographical origin of trait data matters in trait-based analyses. These findings corroborate the assumption that regionally measured trait data and building regional databases are essential for reliable regional-scale trait-based studies. As the significant differences we found for seed mass, specific leaf area and leaf dry matter content most probably resulted from climatic differences between Northwestern Europe and the Pannonian Ecoregion, it is highly advisable that studies analysing these traits in the Pannonian Ecoregion (and possibly in the eastern part of Europe in general) use PADAPT for these traits instead of the LEDA Traitbase or other databases compiling data in regions with markedly different climatic conditions.

## Acknowledgements

We are grateful for all the colleagues who participated in the creation of PADAPT, as well as to everybody who contributed to the LEDA Traitbase. The authors were supported by the National Research, Development and Innovation Office [PT: KKP 144068, K 137573; JS: PD 137747] during the manuscript preparation. The work of JS was supported by the Bolyai János Scholarship of the Hungarian Academy of Sciences [BO/00587/23/8].

